# The Cardioprotective Role of Neutrophil-Specific STING in Myocardial Ischemia/Reperfusion Injury

**DOI:** 10.1101/2024.09.06.611551

**Authors:** Maegan L. Brockman, Triniti A. Scruggs, Lanfang Wang, Gabriella Kabboul, John W. Calvert, Rebecca D. Levit

## Abstract

**Background:** Neutrophils are the most rapid and abundant immune cells to infiltrate the myocardium following myocardial ischemia/reperfusion injury (MI/R). Neutrophil heterogeneity has not been well characterized in MI/R, and studies have shown conflicting results regarding the impact of neutrophil depletion on cardiac injury. We thus aim to study the impact of neutrophils with enriched type I interferon signature and the role of STING (stimulator of interferon genes) signaling in neutrophils on cardiac reperfusion injury.

**Methods:** We utilized single-cell RNA sequencing to study neutrophil heterogeneity in response to MI/R. We generated a neutrophil-specific STING knockout mouse to assess the role of neutrophil STING in a model of MI/R. We examined cardiac function following injury via echocardiography and assessed the immune cell trajectory following injury utilizing flow cytometry.

**Results:** We identified a population of neutrophils with enriched type I interferon signaling and response to type I interferon following MI/R. We found that genetic deletion of neutrophil-specific STING led to worsened cardiac function following MI/R. Further investigation of the immune response by flow cytometry revealed decreased neutrophil infiltration into the myocardium and a shift in macrophage polarization.

**Conclusions:** Our findings suggest that neutrophil-specific STING is cardioprotective in MI/R, partly due to its effects on downstream immune cells. These results demonstrate that early alterations or therapeutic interventions can influence key events in the resolution of inflammation following MI/R.

## Introduction

Ischemic heart disease is currently the leading cause of death worldwide (1, 2). Myocardial infarction (MI) occurs when blood flow to the myocardium is blocked, leading to cardiomyocyte death (1, 2). Timely restoration of blood flow, or reperfusion, is the current standard of care for patients; however, reperfusion can lead to death of previously viable cardiomyocytes, termed reperfusion injury (3). In myocardial ischemia-reperfusion injury (MI/R), inflammation plays a significant role in damaging cardiomyocytes and influencing infarct size, with neutrophils being the most rapid and numerous cells to infiltrate the heart within the first 24 hours (4, 5). While neutrophils play an essential role in wound healing, they can cause further damage through the generation of reactive oxygen species (ROS), proteolytic enzymes, and neutrophil extracellular traps (NETs) (6). Multiple studies have found neutrophil depletion is cardioprotective after injury (7, 8). In contrast, several studies have highlighted beneficial functions of neutrophils, such as the clearance of dead cells and matrix debris, as well as their influence on the phenotype of macrophages to favor repair and remodeling (9, 10). Neutrophil heterogeneity may explain some of the conflicting findings in these studies.

Neutrophil populations with unique properties have been described in autoimmune diseases, cancer, infection, and stroke, including populations displaying pro- or anti-inflammatory characteristics (11–14). In two published studies of neutrophil heterogeneity after permanent ligation of the coronary artery in mice, there was consistency in the identified neutrophil populations. Both groups identified a pro-inflammatory ‘aged’ neutrophil phenotype identified by the glycoprotein SiglecF, but the contribution of this population to cardiac inflammation has not been experimentally characterized (15–17). These studies also identified a type I interferon (IFN) responding neutrophil population characterized by elevated interferon stimulated genes (15–18). The significance of type I IFN responding neutrophils has not been thoroughly investigated, and many studies investigating type I IFN signaling in MI have utilized global knockouts or inhibitors to target this pathway (15, 19). Since the role of type I IFN signaling in neutrophils is not well known, we sought to investigate the effects of type I IFN signaling in neutrophils in MI/R.

Various inflammatory mediators can polarize neutrophils to either a pro- or anti-inflammatory phenotype, with the most well studied being TNFα, IL-6, and IL-1. The type I IFNs, IFNα and IFNβ are more commonly associated with other immune cell types, but are known to promote the shift of neutrophils to a pro-inflammatory state in diseases such as COVID-19, systemic lupus erythematosus, and cancer (21–24). A major driver of transcription of type I IFNs is the DNA sensor cyclic GMP-AMP synthase (cGAS) and its downstream signaling effector, stimulator of interferon genes (STING) (25). In the heart, neutrophils recruited to the infarcted area may detect extracellular DNA from damaged cells via cGAS-STING signaling, leading to type I IFN production (15, 26). Type I IFNs are cytokines that modulate inflammation by stimulating chemokine and cytokine production, enhancing the expression of interferon-stimulated genes (ISGs), and boosting immune cell functions (27). cGAS/STING activation initiates the phosphorylation and translocation of the transcription factor IRF3 (pIRF3) which drives type I IFN transcription. These IFNs bind to their receptor, activating the JAK-STAT pathway and further transcription of ISGs. ISG-encoded proteins protect host cells from viral infections by inhibiting various stages of the viral lifecycle, including entry, replication, translation, assembly, and release (28).

Clinical trials of immune-modulatory therapies in cardiac ischemia have primarily targeted cytokines or their receptors. The net effects of perturbing these systems are difficult to predict due to the involvement of many immune cell types and processes that utilize these signaling networks, as well as temporal and clinical heterogeneity. Therapies in clinical trials have shown improvement in outcomes after ischemic injury, but benefits have been blunted by risk of infection (29). For example, the anti-inflammatory drugs colchicine and the IL-1β antagonist, canakinumab, have failed to demonstrate overall net clinical benefits for MI, though patients with elevated inflammatory markers may experience greater benefit (30–32). Colchicine did not show improvements in hard endpoints of death and MI, and canakinumab reduced composite events, but increased the risk of fatal infections with no net survival benefit (30–32). There are currently no therapeutics that mitigate acute immune mediated reperfusion injury to improve cardiovascular outcomes.

Given that ischemic heart disease continues to be a major cause of morbidity and mortality, we conducted this study to explore the role of neutrophils with an enriched type I IFN signature in MI/R and to guide future neutrophil-targeted therapies. Our findings revealed that STING-expressing neutrophils have a cardioprotective effect by both modulating neutrophil infiltration into the heart and promoting the polarization of macrophages toward pro-reparative phenotypes. Exploiting the beneficial phenotypes of neutrophils may be therapeutically valuable in MI/R, as it could help minimize harmful functions while preserving reparative processes to reduce cardiac injury. These insights could enhance the development of cardiac-targeted immune therapies during the peri-infarct period.

## Methods

### Animals

To assess the role of neutrophil-specific STING in MI/R, we generated a neutrophil-specific STING KO mouse line for use in experiments. We utilized the Catchup mouse strain (gifted from Dr. Matthew Long of the Ohio State University) expressing Cre recombinase under the Ly6G promoter, which has been shown to be specific to murine neutrophils (Supplemental Fig. 1 A-B) (33). These mice were crossed with mice having *loxP* sites flanking the STING allele (STING^flox^; Jackson 031670) to generate the neutrophil-specific STING KO mice. Neutrophil-specific STING KO mice and aged-matched wild-type littermate controls (C57BL/6J; male; 10-12 weeks of age) were used in all experiments. To establish a well-controlled experimental system, male mice were chosen for this study due to the differences in injury severity between male and female mice in MI/R (34). Animals were housed in the vivarium contained within Emory’s Division of Animal Resources with a 12-h light–dark cycle and access to water and food *ad libitum*. All animal work was approved by the Emory University Institutional Animal Care and Use Committee (IACUC).

### Genotyping

Ear biopsies of mice 4-6 weeks old were incubated at 55 °C for overnight in DirectPCR lysis buffer (Viagen 402-E) containing 0.3 mg/ml proteinase K (Thermo AM2546). The enzyme was heat inactivated by incubating samples at 85° for one hour after digestion. Primers used for the detection of the wild-type Ly6G allele were: Forward 5′-GGTTTTATCTGTGCAGCCC-3′; Reverse 5′-GAGGTCCAAGAGACTTTCTGG-3′) and for the detection of the mutant Ly6G allele were: Forward 5′-ACGTCCAGACACAGCATAGG-3′; Reverse 5′-GAGGTCCAAGAGACTTTCTGG-3′). Primers used for the detection of the floxed STING allele were: Forward 5’-TTTTCATCTGCCTTCCAGGT-3’; Reverse 5’-GCGCACACACACTAAAAACTG-3’. Primers and DNA samples were added to a PCR Master Mix (Thermo K0171). The following PCR conditions were applied: 1 min, 95 °C initial denaturation; 15 sec, 95 °C cyclic denaturation; 15 sec, 60 °C cyclic annealing; 35 sec, 72 °C cyclic elongation for a total of 35 cycles, followed by a 10-min 72 °C elongation step. PCR amplification products were analyzed by agarose gel electrophoresis.

### Myocardial Ischemia Reperfusion Model

Mice 10-12 weeks of age were anesthetized with intraperitoneal injections of ketamine (80-100 mg/kg) and xylazine (5-10 mg/kg), intubated, and connected to a rodent ventilator. Prior to the surgical procedure, mice also received an intraperitoneal injection of 200U/kg sodium heparin. A median sternotomy was performed, and the proximal left coronary artery was ligated with a 7-0 silk suture for a duration of 60 minutes. A short segment of PE-10 tubing was placed between the LCA and 7-0 silk suture to minimize damage and allow for complete reperfusion following the ischemic period. After 60 minutes, the ligature and tubing were removed, and reperfusion was confirmed. The chest wall and skin incision were closed with 4-0 Monocryl suture and a monofilament suture respectively. Mice were given buprenorphine extended-release (0.5mg/kg) as an analgesic after surgery and 72 hours post-surgery if needed.

### RNA Sequencing

Cells were isolated from the left ventricle following MI/R (see detailed section below), and neutrophils were enriched via negative immunomagnetic selection (STEMCELL 19762). For our studies gene expression libraries were sequenced using an Illumina NovaSeq600 targeting a depth of 50,000 reads per cell in collaboration with the Emory Primate Center Genomics Core Laboratory. Cell Ranger software (10x Genomics) was used for low-level analysis, including demultiplexing, mapping to a reference transcriptome, and annotation of unique molecular identifiers. All subsequent scRNA-seq analysis was performed using the Seurat R package (v4.0.1) pipeline.

### Gene Expression Analysis of Cardiac Interferons and Interferon Stimulated Genes

Quantitative real-time PCR (qRT-PCR) was performed to measure the expression levels of the type I interferons and interferon stimulated genes in cardiac tissue. RNA was isolated using RNeasy Kits for RNA Purification (Qiagen 74106). Synthesis of cDNA was performed using High-Capacity cDNA Reverse Transcription Kit’s (Thermo Fisher Scientific 4374966) protocol. The housekeeping gene (*Gapdh* Taqman: Mm99999915) was used as an endogenous control and all genes expressed were normalized to the *Gapdh* housekeeping gene using the ddCT method. The primers used are as follows: *Ifna1* TaqMan: Mm03030145; *Ifnb1* TaqMan: Mm00439552; *Ifit2* TaqMan: Mm00492606; *Isg15* TaqMan: Mm01705338; *Cxcl10* TaqMan: Mm00445235; *Tlr9* TaqMan: Mm00446193.

### Isolation of Immune Cells from the Heart and Blood

Leukocytes were isolated from the left ventricle (LV) following MI/R. Mice were anesthetized with 4% isoflurane and blood was obtained via the inferior vena cava and transferred into EDTA tubes. The heart was extracted and perfused with 10 ml of ice-cold phosphate-buffered saline (PBS) to exclude circulating blood cells. The atria and free right ventricular wall were removed and the LV was dissected and minced with fine scissors. The tissue was enzymatically digested in 800 µl of solution containing 450U/ml collagenase I (from *Clostridium histolyticum;* Sigma C0130), 60 U/ml Deoxyribonuclease I (DNase I) (from bovine pancreas; Sigma D4513), and 60 U/ml hyaluronidase (from bovine testes; Sigma H3506) dissolved in Dulbecco’s Modified Eagle’s Medium (DMEM) (Sigma D5671) for 60 minutes at 37°C under gentle agitation. Following digestion, cellular digest was kept at 4°C for the remainder of processing. The digestion was passed through a 40-µm cell strainer to exclude cardiomyocytes and debris and centrifuged at 400 x g for 5 minutes. The cell pellet was resuspended in 1 ml of red blood cell (RBC) lysis buffer (Sigma R7757) and incubated at room temperature for 5 minutes. Lysis was stopped by adding 10 ml of cold Hank’s balanced salt solution (HBSS) and cells were centrifuged at 400 x g for 5 minutes. Blood leukocytes were isolated by adding 10 ml of prewarmed RBC lysis buffer to blood and the mixture was incubated at room temperature for 10 minutes. Lysis was stopped by adding 20ml of cold HBSS and cells were pelleted at 400x g for 5 minutes. Cells were resuspended in FACs buffer for flow cytometric analysis.

### Flow Cytometric Staining of Immune Cells

Isolated cells were resuspended in FACs buffer (PBS, 0.02% Fetal Bovine Serum (FBS), and 1mM EDTA at a density of 3 x 10^5^ cells. Cells were resuspended in Fc blocking buffer (BD 553142). After blocking, cells were spun down and resuspended in the primary antibody staining buffer, including viability dye to gate out dead cells (eBioscience 65-0866-18). Primary antibodies used for staining are listed in Table 1. Cells were fixed and permeabilized using either BD Cytofix/Cytoperm™ Fixation/Permeabilization Solution Kit (BD Biosciences 554714) or Biolegend True-Nuclear™ Transcription Factor Buffer Set (Biolegend 424401) for cytokine or transcription factor staining. Intracellular staining was performed after permeabilization in permeabilization buffer. Cells were washed and resuspended for flow cytometry analysis. Cells were acquired on a Cytek Aurora™ and analyzed using FlowJo 10.

**Table 1:** Flow Cytometry Antibodies.

| *Marker* | *Fluorophore* | *Catalog Number* | *Dilution* |
| --- | --- | --- | --- |
| CCR2 | Brilliant Violet 421 | Biolegend 150605 | 1:100 |
| CD11b | PerCP | Biolegend 101229 | 1:100 |
| CD45 | Alexa Fluor 700 | Biolegend 103127 | 1:200 |
| CD64 | Brilliant Violet 711 | Biolegend 139311 | 1:50 |
| CD206 | PE-Fire 700 | Biolegend 141741 | 1:100 |
| CXCL10 | Alexa Fluor 594 | Bioss bs-1502R-594 | 1:100 |
| IFNα | FITC | Pbl assays 22100-3 | 1:100 |
| Ly6G | APC-Fire 750 | Biolegend 127651 | 1:50 |
| pIRF3 | APC | Cell Signaling 81093 | 1:50 |
| TLR9 | PE | Biolegend 159103 | 1:50 |

### Echocardiographic Analysis

Mice were anesthetized with isoflurane in 100% O_2_ (5% for induction and 1-1.5% for maintenance) and secured to the Visualsonics Rail System III with medical tape. Core body temperature, respiration rate, and EKG were continuously monitored throughout the procedure. Transthoracic echocardiography of the left ventricle using a scan head interfaced with a Vevo echocardiography system (Visualsonics) was used to obtain high resolution B-mode and M-mode images.

### Histology (Masson’s Trichrome Staining)

Tissues were fixed overnight in 10% formalin, routinely processed, paraffin-embedded, and sectioned at 5µM. Paraffin embedded sections were subjected to deparaffinization in xylene, rehydrated in a graded series of ethanol, rinsed with deionized water, and fixed in Bouin’s Solution (Sigma HT10132) at room temperature overnight. Slides were then rinsed in a container with running tap water until clear, then rinsed once in diH2O and stained with Weigert’s Iron Hematoxylin solution (Sigma 1159730002) for 5 minutes. Slides were rinsed in tap water and then diH2O and stained with Biebrich Scarlet-Acid Fuschin (Sigma HT151) for 5 minutes. Rinsing in tap water then diH2O was repeated, followed by 5 min in 5% phosphotungstic/phosphomolybdic acid solution for 5 min (Sigma HT152 and HT153 respectively). Slides were rinsed in diH2O then stained for 4 min in Aniline Blue solution, then rinsed again in diH2O, and set in 1% acetic acid for 2 min. Slides were then dehydrated by consecutive two dips in diH2O, 75%, 95%, and 100% ethanol, then twice in xylene, and mounted using Cytoseal (Richard-Allan Scientific 8310–4). Images were taken at 40x with a NanoZoomer SQ Whole Slide Scanner (Hamamatsu) and analyzed using ImageJ software.

### Western Blot

Neutrophils isolated from the peritoneum of WT and STING KO mice were collected and lysed in RIPA buffer containing protease inhibitors. Protein concentrations were measured with the Pierce™ BCA Protein Assay Kits (Thermo Scientific 23209). Equal amounts of protein were loaded into lanes of 12% SDS-PAGE gels. The protein was then transferred to a polyvinylidene difluoride membrane. The membranes were then blocked and probed with primary antibodies at 1:1000 dilution (STING (D1V5L): Cell Signaling 50494; H3 (1B1B2): Cell Signaling 14269) overnight at 4°C. Immunoblots were next processed with secondary antibodies for 1 hour at room temperature. Immunoblots were then probed with ECL Western Blotting Substrate (Thermo Scientific PI32209) to visualize signal, followed by visualization using film. Data were analyzed using ImageJ. Protein was normalized to the housekeeping gene, H3.

### Statistics

Statistical analysis was performed on GraphPad Prism 10. Unless otherwise stated, values are plotted as mean +/− standard deviation. Statistical significance was evaluated as follows: (1) unpaired Student *t* test for comparison between 2 means; (2) a 1-way ANOVA with a Bonferroni test as the post hoc analysis for comparison among ≥3 groups; and (3) a 2-way ANOVA with a Bonferroni test as the post hoc analysis for comparison among the means from groups of wild-type (WT) and STING knockout mice. For nonparametric tests, Mann-Whitney U test was used for comparison of 2 variables. A *p*-value <0.05 was considered significant.

## Results

### Single-cell RNA Sequencing of Neutrophils Post-MI/R Reveals a Type I Interferon Responding Neutrophil Population

Published single cell datasets focused on neutrophils isolated from hearts that underwent ischemia without reperfusion, which has a unique inflammatory and cell injury pattern compared to MI/R (34, 35). To define the heterogeneity of neutrophils after MI/R we performed single-cell RNA sequencing on cardiac-infiltrating neutrophils isolated from the hearts of mice 24 hours post-MI/R. C57BL/6J mice underwent either 60 minutes of ischemia followed by 24 hours of reperfusion or sham thoracotomy. To capture the largest diversity of neutrophils, cells were enriched by negative immunomagnetic selection for single-cell RNA sequencing (Fig. 1A). Cells from sham (n=1) and MI/R hearts (n=3) were clustered at low resolution to identify major cell types. t-distributed stochastic neighbor embedding (t-sne) was used to visualize the data in 2D space (Fig. 1B). Neutrophils were identified by the expression of canonical neutrophil markers (*S100a8, S100a9, Csf3r, Cxcr2, Mmp9, Csf1*, and *Il1r2*) as described previously (Supplemental Fig. 2) (16). Neutrophils were then subset from the original unintegrated counts matrices, re-scaled, and re-clustered to produce a harmonized data set consisting of 16,165 individual neutrophils (569 from sham and 15,596 from MI/R) in which a median of 1987 expressed transcripts per cell were detected. Unsupervised clustering identified 5 distinct neutrophil populations (Fig. 1C). The majority of neutrophils (53%; 3% of all sham and 54% of all MI/R) segregated to Cluster 1 which was defined by genes associated with cytokine-mediated signaling (Fig. 1 D-E). Cluster 2 consisted of 25% of all neutrophils (11% of all sham and 25% of all MI/R) and was defined by genes associated with neutrophil activation and immunity. Cluster 3 consisted of 8% of all neutrophils (2% of all sham and 8% of all MI/R) and was defined by genes associated with type I IFN signaling and cellular responses to type I IFN (Fig. 1F). Cluster 4 accounted for 10% of all neutrophils and was defined by genes associated with translation. Interestingly, this cluster contained 84% of the sham neutrophils to only 8% of the MI/R neutrophils. Finally, Cluster 5 contained only MI/R neutrophils (5%) and was defined by genes associated with interleukin-12 mediated signaling. Comparing transcriptomes of sham hearts and MI/R-injured hearts by scoring cells for the expression of ISG genes shows significantly upregulated ISG signaling in MI/R hearts (Fig. 1G).

**Figure 1.**
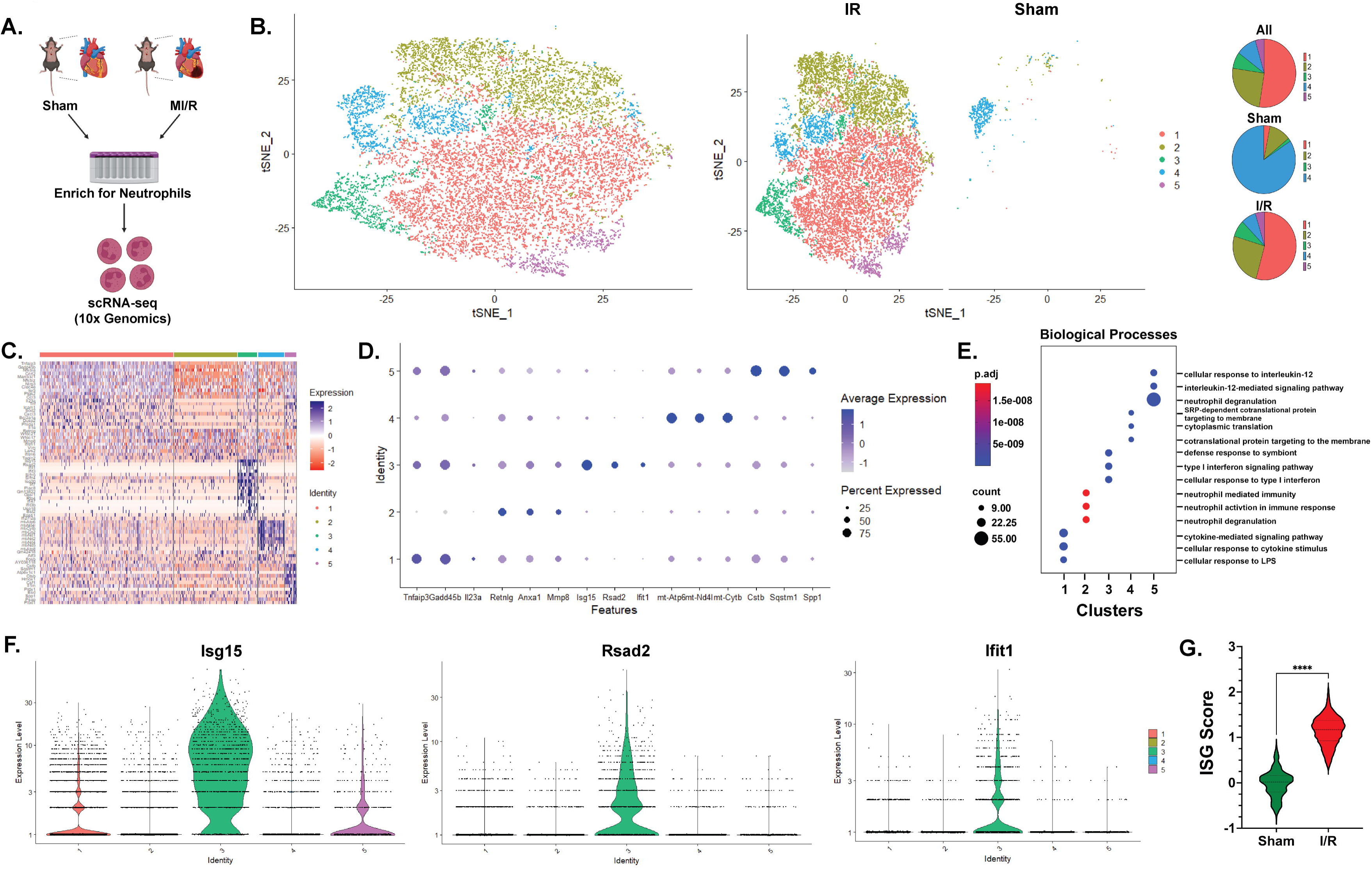
Single-cell RNA Sequencing of Neutrophils Post-MI/R Reveals Type I Interferon Responding Neutrophil Population: Schematic of experimental design (A). t-SNE plots of sham and MI/R integrated (left) and segregated (right) neutrophils colored by clusters (B). Heat map plotting the top marker genes for each cluster (C). Dot plot of the expression of specific marker genes used to identify neutrophil subtypes (D). Gene set enrichment analysis of MI/R neutrophil clusters (E). Violin plots showing the expression of representative interferon stimulated genes in each neutrophil cluster (F). Violin plot of aggregate ISG score in sham and MI/R neutrophils (G). ***p<0.001, Mann-Whitney test.

### Type I Interferons and Interferon Signaling are Present in Neutrophils Early Post-MI/R

To validate single-cell RNA sequencing results, we next collected cardiac-tissue 24 hours after MI/R and evaluated the type I IFN response. Male C57BL/6J mice underwent either 60 minutes of ischemia followed by 24 hours of reperfusion or the sham procedure and cardiac tissue was collected for flow cytometry or gene expression analysis by qPCR. As consistent with previous literature, there was a significant increase in neutrophils present in the heart 24 hours post-MI/R compared to sham hearts (Fig. 2A). Gene expression analysis revealed significantly elevated expression of *Ifna1* and *Ifnb1* (Fig. 2B), indicating the production of type I IFNs is increased in cardiac tissue 24 hours post-MI/R. To assess downstream type I IFN signaling in the heart after MI/R, gene expression of ISGs was assessed in cardiac tissue. Expression of *Ifit2, Isg15, CXCL10,* and *TLR9* were significantly increased in the heart 24 hours post-MI/R (Fig. 2C). Since neutrophils are the most abundant immune cells to infiltrate the heart within the first 24 hours, we wanted to assess type I IFN signaling specifically in neutrophils. We isolated immune cells from the heart following MI/R and stained for markers of type I IFN signaling. Cells were gated on singlets and live cells, and neutrophils were defined as cells that were CD45+, Ly6G+ cells. Flow cytometry analysis shows significant increase in IFNα+ and pIRF3+ neutrophils (Fig. 2D), indicating the activation of the transcription factor IRF3 that initiates transcription of type I IFNs. We also identified significantly increased CXCL10+ and TLR9+ neutrophils in MI/R hearts (Fig. 2D), suggesting that type I IFN signaling is activated in neutrophils.

**Figure 2.**
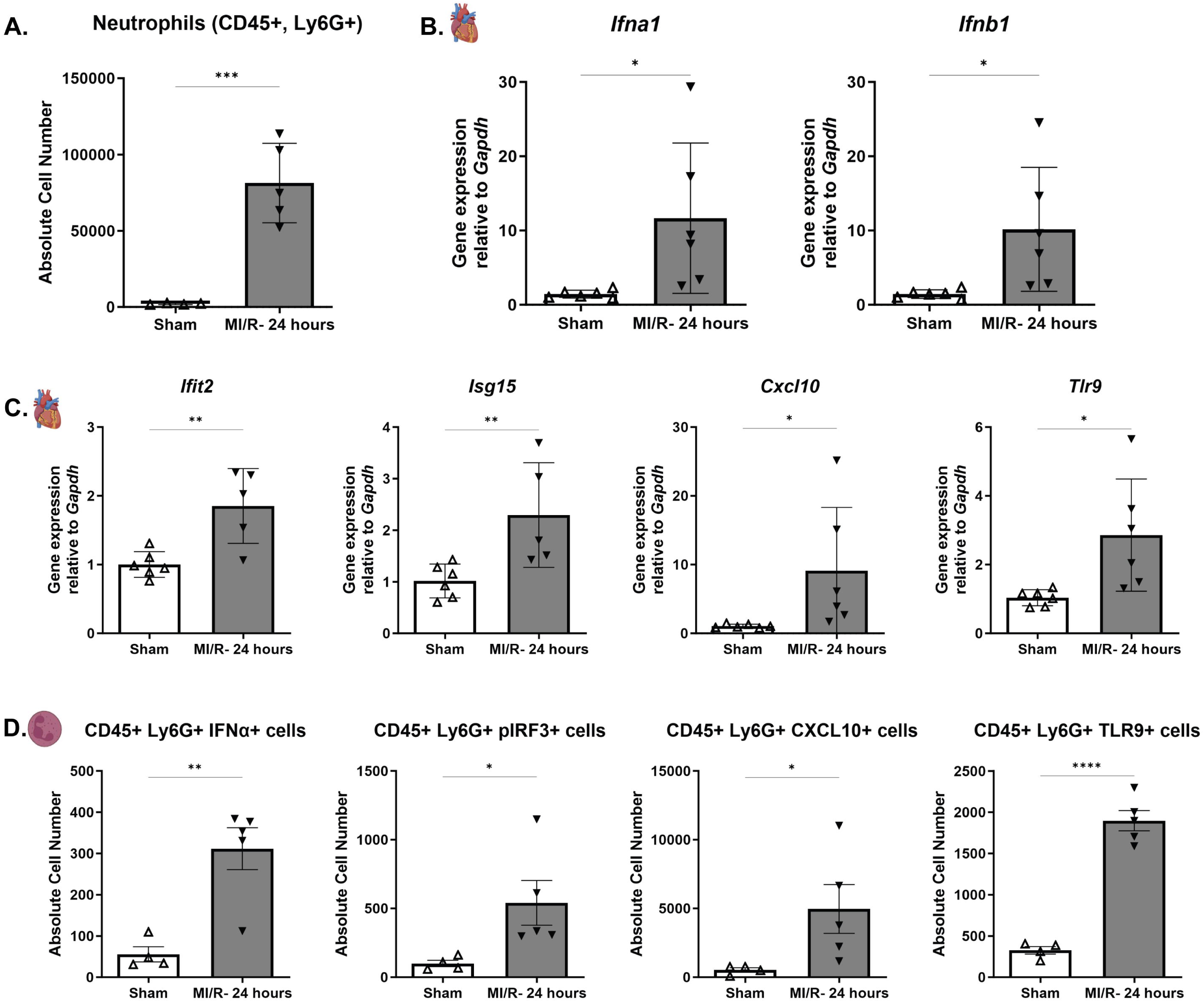
Type I IFN Signaling Early Post-MI/R: Flow cytometry analysis of neutrophils isolated from cardiac tissue of sham and IR mice 1 day after reperfusion (A). Values are mean ± SD. Unpaired one-tailed t test. N=4-5. Gene expression levels of Ifna1 and Ifnb1 in cardiac tissue collected from sham and IR mice 1 day after reperfusion (B). Values are mean ± SD. Unpaired one-tailed t test. N=6. Gene expression levels of interferon stimulated genes in cardiac tissue collected from sham and IR mice 1 day after reperfusion (C). Values are mean ± SD. Unpaired one-tailed t test. N=5-6. Flow cytometry analysis of type I IFN signaling in neutrophils isolated from cardiac tissue of sham and IR mice 1 day after reperfusion (D). Values are mean ± SD. Unpaired one-tailed t test. N=4-5. *p<0.05; **p<0.01; ***p<0.001; ****p<0.0001.

### Knockout of STING in Neutrophils Worsens Cardiac Injury

After identifying a population of neutrophils with enriched type I IFN signaling and further validating the presence of this neutrophil population in MI/R, we generated a neutrophil-specific STING KO mouse to further investigate this signaling pathway in neutrophils in MI/R. There was no significant difference in peripheral blood counts in the neutrophil-specific STING KO mice compared to WT littermate controls (Supplemental Fig. 1C). To validate our knockout efficiency, immune cells were isolated from the peritoneum of WT or neutrophil-specific STING KO mice and the neutrophil fraction was purified by density gradient centrifugation. The fraction of non-neutrophil cells (macrophages, monocytes, and lymphocytes) was collected for analysis to ensure STING protein levels are not affected in other immune cell types. Analysis by Western Blot showed significant reduction of STING protein in neutrophils, but no change in STING protein levels in other non-neutrophil immune cells (Fig. 3A-B). To investigate the effects of neutrophil-specific STING in a model of MI/R, WT or neutrophil-specific STING KO mice were subjected to 60 minutes of ischemia followed by 2 weeks of reperfusion. Cardiac function was assessed by echocardiography at baseline and 2 weeks following reperfusion. Neutrophil-specific STING KO mice showed significantly increased left-ventricular end systolic diameter (LVESD) and left-ventricular end diastolic diameter (LVEDD) compared to WT control mice 2 weeks post-MI/R (Fig. 3C-D). Neutrophil-specific STING KO mice also showed significantly decreased ejection fraction and fractional shortening compared to WT control mice (Fig. 3E), indicating that neutrophil-specific STING KO mice have worse cardiac function following MI/R. Neutrophil-specific STING KO mice also had increased ventricular weight to tibia length (VW/TL) ratios compared to WT mice, and Masson’s Trichrome staining for fibrosis revealed significantly increased fibrosis of the left ventricle in neutrophil-specific STING KO mice (Fig. 3F-H). These results suggest that neutrophil-specific STING is important for healing in MI/R and loss of STING in neutrophils leads to worse cardiac function and increased fibrosis.

**Figure 3.**
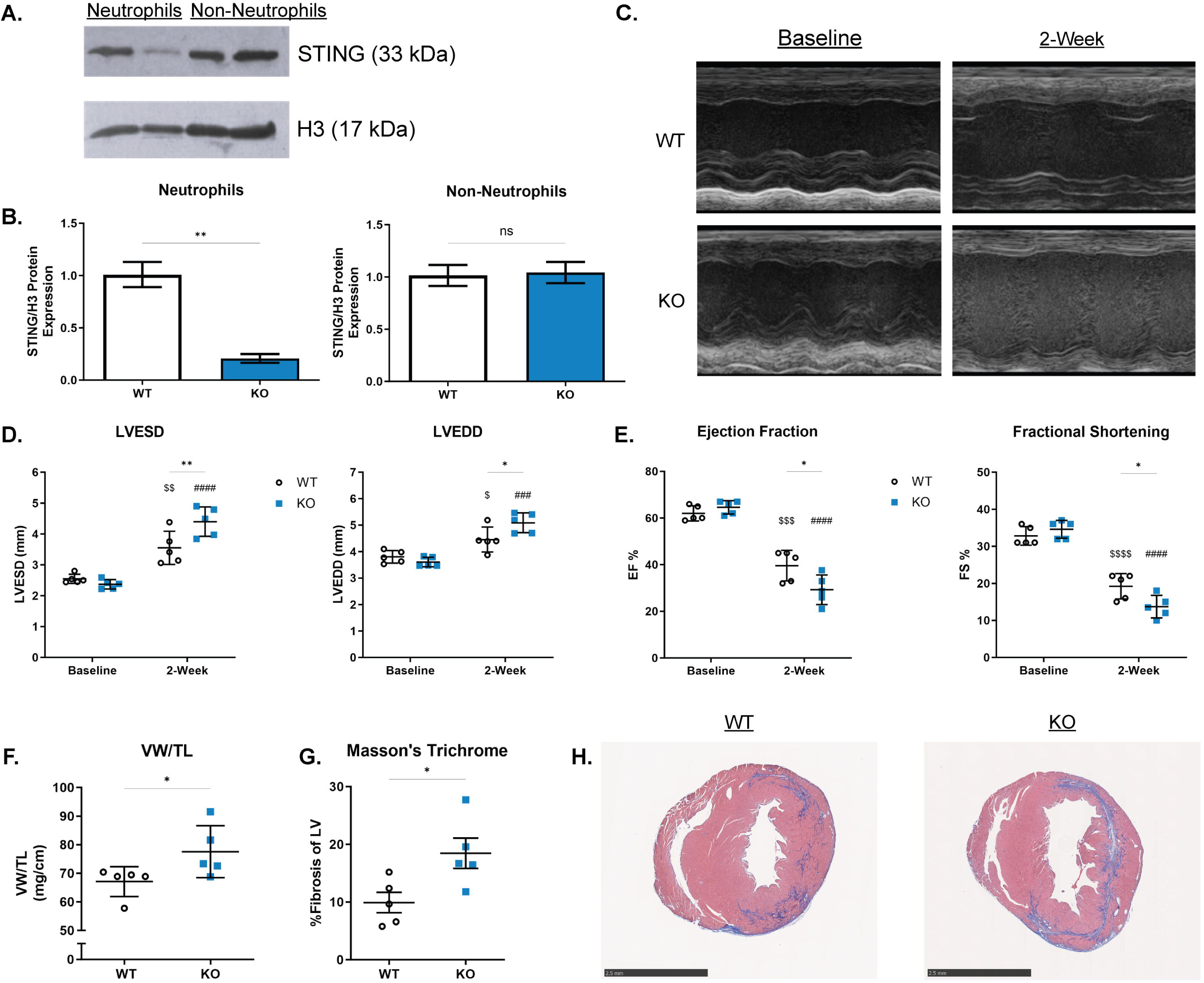
Knockout of STING in Neutrophils Worsens Cardiac Injury: Representative immunoblot **(A)** and quantification of STING knockout efficiency in neutrophils and non-neutrophils **(B).** Values are mean ± SD. Unpaired two-tailed t test. N=3. Representative echocardiography images of parasternal long-axis left ventricular M-mode **(C).** Left ventricular end-systolic diameter, left ventricular end-diastolic diameter, **(D)** left ventricular ejection fraction, and fractional shortening **(E)** from wild-type and neutrophil-specific STING KO mice at baseline and two weeks after ischemia/reperfusion. Values are mean ± SD. Two-way RM ANOVA with Bonferroni. ^$^p<0.0.05; ^$$^p<0.01; ^$$$^p<0.001; ^$$$$^p<0.0001. vs WT baseline. ^#^p<0.0.05; ^##^p<0.01; ^###^p<0.001; ^####^p<0.0001. vs KO baseline. N=5. Ratios of ventricular weight to tibia length were used as a measure of cardiac hypertrophy **(F)** and Masson’s Trichrome was used to stain for fibrosis and quantified two weeks after ischemia/reperfusion **(G,H).** Scale bare equals 2.5 mm. Values are mean ± SD. Unpaired one-tailed t test. N=5. LV = left ventricle; LVESD = left ventricular end systolic diameter; LVEDD = left ventricular end diastolic diameter; EF = ejection fraction; FS = fractional shortening; VW/TL = ventricular weight/tibia length. *p<0.0.05; **p<0.01; ***p<0.001; ****p<0.0001.

### Knockout of STING in Neutrophils Alters Immune Response in MI/R

To further investigate the role of STING in MI/R, we assessed the immune response in WT and neutrophil-specific STING KO mice following MI/R. WT or neutrophil-specific STING KO mice were subjected to 60 minutes of ischemia followed 24- or 72-hours of reperfusion or the sham thoracotomy, and immune cells were isolated from the left ventricle for flow cytometry analysis. We found significantly fewer total leukocytes (CD45+ cells) and myeloid cells (classified as CD45+, CD11b+) infiltrating into the myocardium in neutrophil-specific STING KO mice compared to WT mice day 1 (D1) after reperfusion (Fig. 4A). We did not see any difference in the number of lymphocytes (CD45+, CD11b-) found in the myocardium of WT and neutrophil-specific STING KO mice (Fig. 4A). The differences in leukocytes and myeloid cells in the myocardium D1 after reperfusion was driven by significantly less neutrophils (CD45+, CD11b+, Ly6G+ cells) in the hearts of neutrophil-specific STING KO mice. (Fig. 4B). There was also a significant decrease in neutrophils in WT hearts from D1 to day 3 (D3), but no significant change in neutrophil numbers in neutrophil-specific STING KO hearts from D1 to D3 (Fig. 4B), suggesting a dysregulated acute immune response.

**Figure 4.**
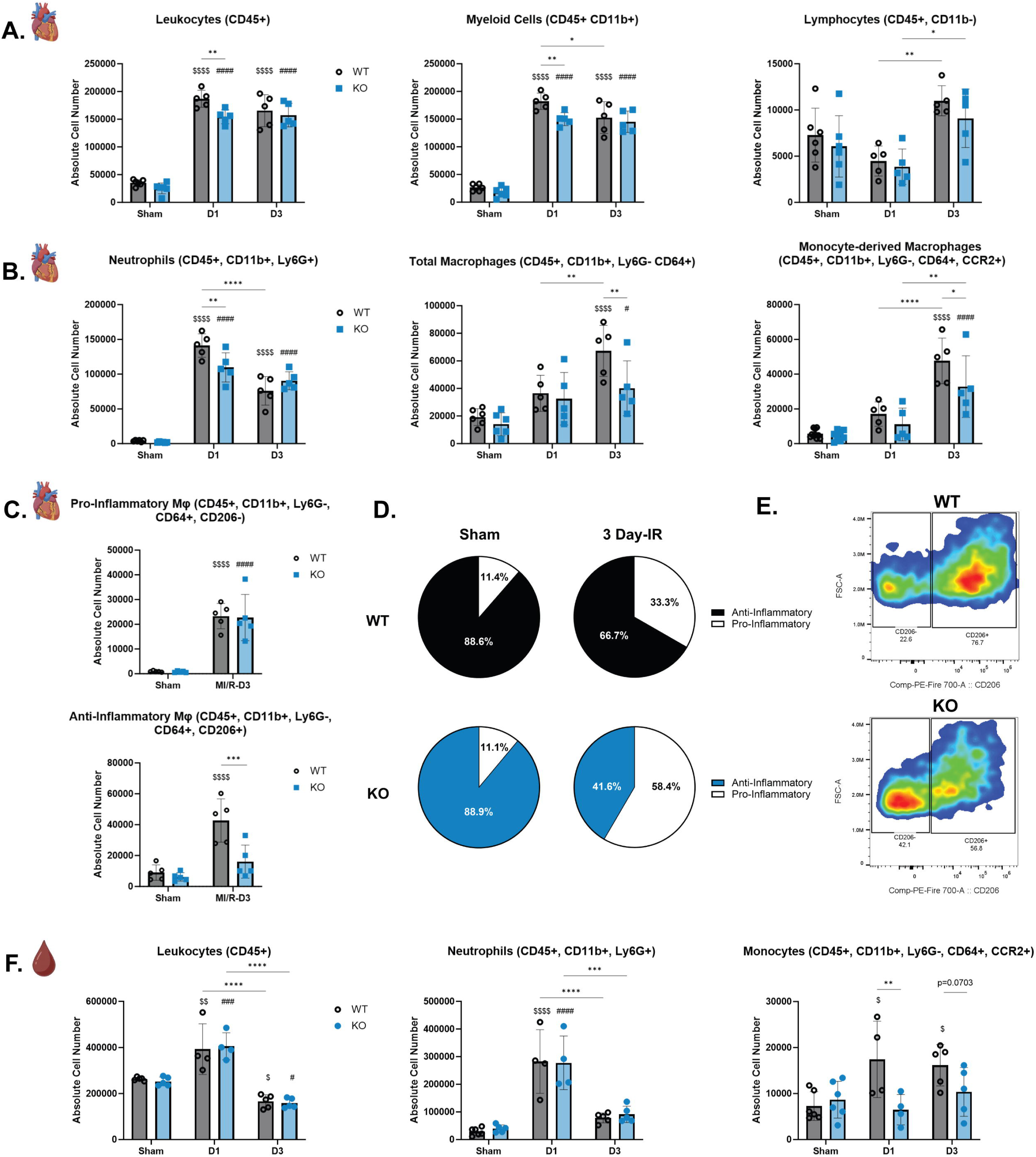
Knockout of STING in Neutrophils alters Immune Response in MI/R: Flow cytometry analysis of total leukocytes, myeloid cells, and lymphocytes isolated from cardiac tissue of sham and IR mice 1- and 3-days after reperfusion in wild-type and neutrophil-specific STING knockout mice (A). Flow cytometry analysis of neutrophils, CD64+ cells, and CCR2+ monocytes isolated from cardiac tissue of sham and IR mice 1- and 3-days after reperfusion in wild-type and neutrophil-specific STING knockout mice (B). Flow cytometry analysis of pro- and anti-inflammatory macrophages isolated from cardiac tissue of sham and IR mice 3-days after reperfusion in wild-type and neutrophil-specific STING knockout mice (C). Percentage of pro- and anti-inflammatory macrophages isolated from cardiac tissue of sham and IR mice 3-days after reperfusion in wild-type and neutrophil-specific STING knockout mice (D). Representative flow cytometry plots of CD206 expression on macrophages isolated from cardiac tissue of wild-type and neutrophil-specific STING KO mice 3-days after reperfusion. (E). Flow cytometry analysis of peripheral blood leukocytes, neutrophils, and monocytes taken from sham and IR mice 1- and 3-days after reperfusion in wild-type and neutrophil-specific STING knockout mice (F). Values are mean ± SD. N=4-6. *p<0.0.05; **p<0.01; ***p<0.001; ****p<0.0001. ^$^p<0.0.05; ^$$^p<0.01; ^$$$^p<0.001; ^$$$$^p<0.0001. vs WT sham. ^#^p<0.0.05; ^##^p<0.01; ^###^p<0.001; ^####^p<0.0001. vs KO sham.

We next investigated if changes in neutrophil infiltration at D1 influenced immune cell number and phenotype later in the healing time course. Three days after reperfusion, the number of CD64+ macrophages found in the heart, which we furthered classified into monocyte-derived macrophages depending on additional phenotypic markers, was significantly less in neutrophil-specific STING KO mice compared to WT mice (Fig. 4B). Analyzing markers for monocytes revealed significantly less monocyte-derived macrophages (CD45+, CD11b+, Ly6G-, CD64+, CCR2+) present in neutrophil-specific STING KO hearts compared to WT hearts D3 after reperfusion (Fig.4B). There was no significant difference in the number of pro-inflammatory macrophages (CD45+, CD11b+, Ly6G-, CD64+, CD206-) found in the hearts of WT and neutrophil-specific STING KO mice, but there was a significant reduction in the number of anti-inflammatory macrophages (CD45+, CD11b+, Ly6G, CD64+, CD206+) found in neutrophil-specific STING KO mice compared to WT mice at D3 (Fig. 4C). Proportionally, we observed a significantly greater percentage of pro-inflammatory macrophages in STING KO mice after MI/R (Fig. 4D-E). Similarly, we saw a significantly lower percentage of anti-inflammatory macrophages in the myocardium of neutrophil-specific STING KO mice compared to WT mice. (Fig. 4D-E). We further investigated whether changes in cardiac-infiltrating immune cells could be due to alterations in circulating cells. We analyzed the number of cells in the blood of WT or neutrophil-specific STING KO mice subjected to 60 minutes of ischemia followed by 24- or 72-hours of reperfusion or the sham thoracotomy (Fig. 4F). There was no significant difference in the number of total circulating leukocytes or neutrophils between WT and neutrophil-specific STING KO mice D1 or D3 after reperfusion; however, we saw significantly fewer circulating monocytes in neutrophil-specific STING KO mice at D1 compared to WT mice (Fig. 4F). These results suggest neutrophil specific STING is important in regulating the acute (D1) neutrophil and later (D3) macrophage response to cardiac injury.

## Discussion

The role of neutrophils in health and disease is shifting away from the simplistic view that they are solely pro-inflammatory and possess a singular phenotype. Instead, there is growing recognition of neutrophil heterogeneity, with a diversity of phenotypes that shape the immune response in both health and disease. Numerous studies have identified a neutrophil population with a type I IFN signature by utilizing single-cell RNA sequencing across varying disease states (13, 36–38). For instance, type I IFN signaling in neutrophils and type I IFN signaling regulating the neutrophil response has been implicated in viral diseases, such as COVID-19, autoimmune diseases, such as systemic lupus erythematosus, and cancer (21–24, 39).The neutrophil type I IFN signature has not been well characterized in MI/R, and we show for the first time that neutrophil-specific STING is important for cardiac healing after reperfusion injury.

Several studies have implicated the cGAS-STING pathway as an important target in MI. These studies largely focus on activation of the cGAS-STING pathway and subsequent type I IFN signaling in cardiomyocytes, macrophages, and monocytes. Studies investigating inhibition of STING with the small molecule inhibitor, H151, showed STING inhibition preserved myocardial function after MI, attenuated cardiac fibrosis, and decreased apoptosis of cardiomyocytes (19, 40). It is reported that treatment with H151 also abrogated the type I IFN response initiated in bone marrow-derived macrophages elicited by dsDNA isolated from hearts (19). Furthermore, inhibition of cGAS in MI promoted the polarization of anti-inflammatory, reparative macrophages, which attenuated adverse cardiac remodeling (41). A complementary study also implicated macrophages as the cell type responsible for phagocytosing DNA and initiating the activation of cGAS-STING signaling to produce type I IFNs (32). This study further investigated inhibition of the cGAS-STING pathway by utilizing global cGAS knockout mice, mutagenized STING mice, and IRF3 knockout mice (15). Interestingly, only cGAS and IRF3 knockout mice showed improvement in cardiac function in an MI model, while the mutagenized STING mice did not show any benefit in survival or preservation of cardiac function post-MI (15). These results suggest a potentially differential role of STING activation in type I IFN signaling as loss of STING activation did not attenuate cardiac injury. These results also underscore the importance of cell-specific and temporal contributions in MI/R, as the loss of neutrophil-specific STING altered the immune cell trajectory and resulted in worsened cardiac function. It is important to emphasize that previous studies have utilized global knockout mice or global inhibitors to target the cGAS-STING-type I IFN pathway in MI. Much like clinical trials of immune modulators, these strategies do not target specific cell types, many of which utilize the same receptors and signaling pathways. Consequently, predicting and understanding the mechanisms behind phenotypic outcomes can be challenging. A more nuanced and targeted therapeutic approach may be needed.

The benefit or harm of neutrophils in myocardial ischemia has also been highly debated. Studies have reported that abrogating the neutrophil response post-MI improves infarct size and mitigating neutrophil recruitment to the myocardium reduces reperfusion injury and recovers cardiac function (7, 8). Conversely, there have been reports that the initial neutrophil response influences many aspects of later inflammatory cell infiltration including cell type and number, and dampening the neutrophil response is detrimental to cardiac healing (10, 42). Several studies in animal models have targeted neutrophils and subsequently saw improvements in cardiac function, decreased scar size, and decreased inflammation. Antagonism of the chemokine receptor CXCR2 attenuated neutrophil infiltration into the myocardium, reduced infarct size, and improved cardiac function in a permanent ligation MI model (43). Inhibition of the alarmin S100A9 during the acute inflammatory phase after MI reduced the number of neutrophils infiltrating into the myocardium and improved cardiac function (44). Metoprolol was shown to inhibit neutrophil migration in an ADRB1-dependent manner and acts to reduce infarct size in MI patients (45). Gasdermin D is required for early mobilization of neutrophils into the area of infarct, and loss of gasdermin D resulted in decreased levels of neutrophils and monocytes in the infarcted heart (46). Additionally, knockout of gasdermin D in mice significantly reduced infarct size, improved cardiac function, and improved overall survival after MI (46). However, there have been many studies demonstrating that abrogating the neutrophil response post-MI can result in worsened outcomes. Extended blockade of S100A9 led to deterioration of cardiac function and increased left ventricle dilation (47). Neutrophil depleted mice subjected to MI had worsened cardiac function, increased fibrosis, and eventually developed heart failure (10). Neutrophils were shown to promote the polarization of macrophages to a reparative phenotype (10). Previous studies have demonstrated the role of netrin-1 in attenuating MI/R, and it was shown that neutrophil-derived netrin-1 attenuated reperfusion injury through myeloid adenosine A2b signaling (48–50). Neutrophils were also shown to be required to induce upregulation of TGF-β1 expression in fibroblasts, which is a key determinant in terminating the pro-inflammatory phase after MI (42). Several studies have also implicated the importance of neutrophil extracellular traps (NETs) in promoting proper cardiac healing post-MI (51, 52). These conflicting results highlight the significance of neutrophil heterogeneity and suggest that varying neutrophil phenotypes may impact different aspects of healing in MI/R.

Many of these studies have also focused on cGAS-STING and type I IFN signaling in models of permanent ligation of the coronary artery. While both permanent ligation of the coronary artery and MI/R involve ischemic insult, the types of injury recapitulated by each model are distinct and the models should not be considered interchangeable (34). Different cellular pathways are activated by ischemia alone versus ischemia followed by reperfusion (34). Thus, activation of cGAS-STING and type I IFN signaling may be differentially affected in each model. Additionally, the immune cell trajectory is distinct in permanent ligation and MI/R (53). In permanent ligation of the coronary artery, total leukocyte numbers peak later after onset of MI, whereas in MI/R total leukocytes peak earlier after onset of MI/R (53). In MI/R, neutrophil infiltration peaks around 24 hours after reperfusion. While the standard of care is to reperfuse patients through percutaneous intervention, not all patients may be reperfused due to various factors (34). Thus, both models are clinically relevant, but results from each model should not be interpreted as the same.

Here we validate the presence of a neutrophil population with enriched type I IFN signaling and response to type I IFN. We demonstrate for the first time that the loss of neutrophil-specific STING negatively impacts cardiovascular outcomes in MI/R, partly by reducing pro-reparative macrophage populations. STING activation in neutrophils is crucial for initiating the immune response, and its loss may impair the downstream cascade necessary for optimal healing following MI/R. Future studies will focus on phenotyping neutrophils in MI/R to determine how STING activation affects the neutrophil response in MI/R, and how changing neutrophil phenotypes caused by loss of STING may affect downstream immune cell responses. We will also assess neutrophil effector functions in neutrophil-specific STING KO mice to determine if loss of STING affects effector functions that contribute to MI/R. Further investigation of the neutrophil phenotype in MI/R and how STING activation affects neutrophil phenotype can help clarify existing contradictions of the harm and benefits of neutrophils after MI/R, and could provide a clinically feasible point of intervention in MI/R to improve cardiovascular outcomes.

## Acknowledgments

Research reported in this publication was supported in part by the Pediatrics/Winship Flow Cytometry Core of Winship Cancer Institute of Emory University, Children’s Healthcare of Atlanta and NIH/NCI under award number P30CA138292. The content is solely the responsibility of the authors and does not necessarily represent the official views of the National Institutes of Health.

## Author Contributions

M.L.B, T.A.S, L.W, G.K, and J.W.C performed experiments and acquired data. M.L.B, J.W.C, and R.D.L interpreted and analyzed data. M.L.B wrote the manuscript and J.W.C and R.D.L revised the manuscript. R.D.L conceived the project.

---

**Supplemental Figure 1.**
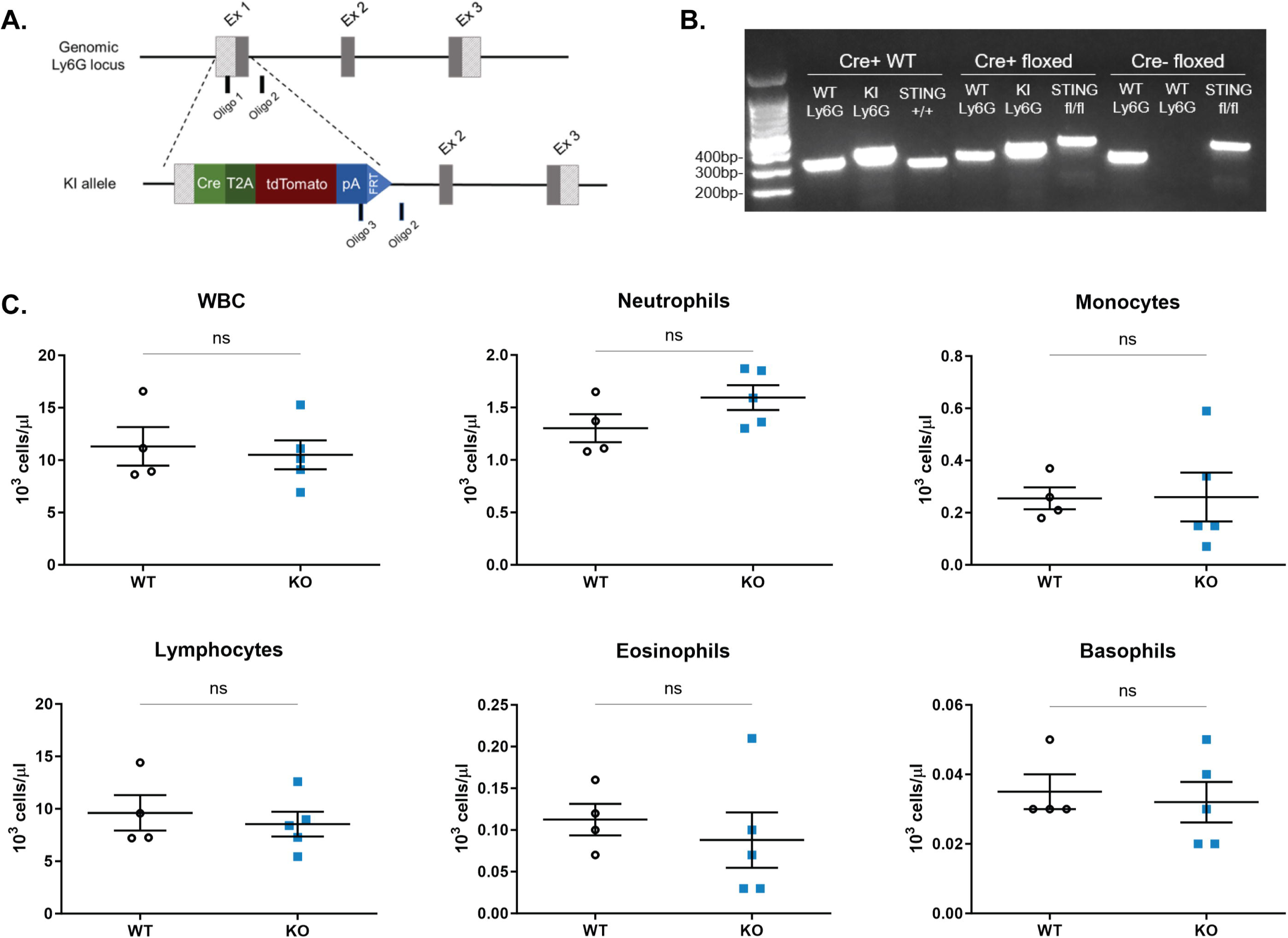
Genotyping of Neutrophil-Specific STING KO Mice: Neutrophil-specific knockout mice were bred by crossing mice heterozygous for Cre recombinase under the Ly6G promoter **(A)** (Hasenberg, Nature Methods, 2015) with mice homozygous for floxed STING purchased from Jackson Labs (#031670). The wildtype Ly6G allele appears at 324bp; the Ly6G knock-in allele appears at 373bp; the wildtype STING allele appears at 298bp and the STING floxed allele appears at 450bp **(B).** CBC analysis of wild-type and neutrophil-specific STING KO mice show no significant differences in cell counts **(C)**.

**Supplemental Figure 2.**
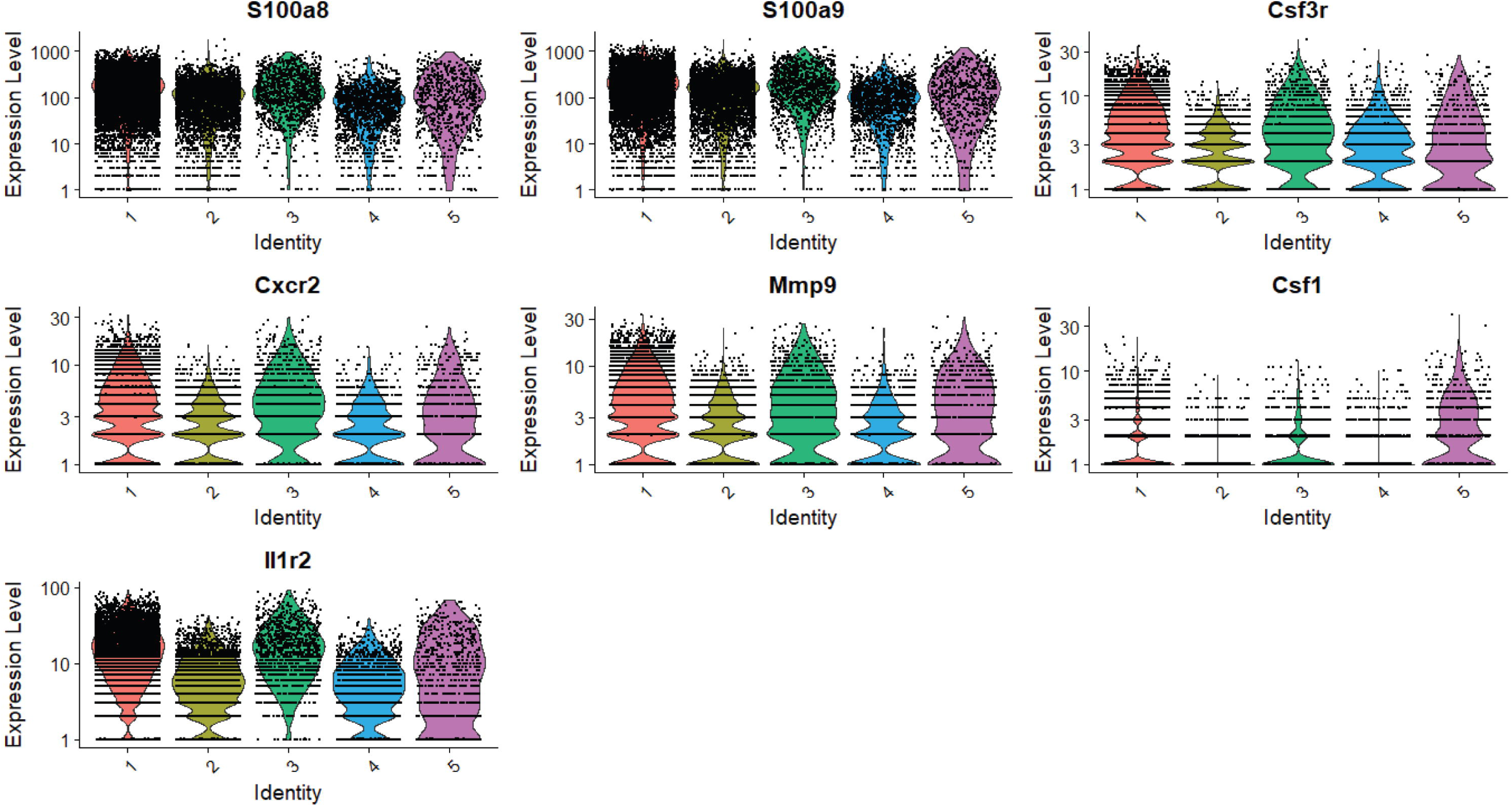
Neutrophil Markers in Single-Cell RNA Sequencing: Violin plots showing the expression of representative genes of neutrophils (including S100a8, S100a9, Csf3r, Cxcr2, Mmp9, Csf1, Il1r2) in each cluster.

